# *Trypanosoma cruzi* PARP is enriched in the nucleolus and is present in a thread connecting nuclei during mitosis

**DOI:** 10.1101/2022.04.07.487537

**Authors:** María Laura Kevorkian, Salomé C. Vilchez Larrea, Silvia H. Fernández Villamil

## Abstract

Poly (ADP-ribose) polymerase (PARP) is responsible for the synthesis of ADP-ribose polymers, which are involved in a wide range of cellular processes such as preservation of genome integrity, DNA damage signaling and repair, molecular switch between distinct cell death pathways and cell cycle progression. Previously, we demonstrated that the only PARP present in *T. cruzi* migrates to the nucleus upon genotoxic stimulus. In this work, we identify the N-terminal domain as being sufficient for TcPARP nuclear localization and describe for the first time that TcPARP is enriched in the parasite nucleolus. We also describe that TcPARP is present in a thread that connects two dividing nuclei and co-localizes with nucleolar material and microtubules. Furthermore, ADP-ribose polymers could also be detected in this wire during mitosis. These findings represent a first approach to new potential TcPARP functions inside the nucleus and will help understand its role well beyond the largely described DNA damage response protein in trypanosomatids.

## Introduction

Poly (ADP-ribose) polymerase (PARP) catalyzes the transfer of ADP-ribose units from NAD^+^ to a target protein or an elongating polymer chain, a post-translational modification known as PARylation (1). Enzymes able to carry out PARylation have been described in all types of eukaryotic organisms, from mammals and plants to lower eukaryotes, except for yeast. Orthologs have also been found in bacteria (2). In humans, the PARP family consists of 18 members grouped into subfamilies. One of these subfamilies, the “DNA-dependent” PARPs (PARP-1, PARP-2 and PARP-3), contain domains capable of recognizing discontinued DNA, thus enabling the recruitment of proteins involved in DNA repairing (3–5). The most extensively studied function of human PARP-1 (hPARP-1) is its role in DNA damage signaling and repair. Nevertheless, over the last years, the functions of poly (ADP-ribose) (PAR) have transcended from being solely associated with genomic damage to being involved in diverse processes such as the regulation of transcription factors, inflammation, cell death and survival, as well as different pathologies (4,6–9).

Our research team has pioneered the study of PAR metabolism in trypanosomatids, specially *Trypanosoma cruzi*, the agent responsible for Chagas’
s disease (10–16). We found a single 65 kDa PARP in *T. cruzi*, named TcPARP, which is expressed in the three developmental stages of the parasite (11). TcPARP shares some structural features with DNA-dependent PARPs, as three of their typical domains are present in the parasite’s counterpart: WGR domain, central regulatory domain and catalytic domain. TcPARP lacks the typical zinc-finger N-terminal PARP-1 domain but bears a shorter basic N-terminal region, like hPARP-2 and hPARP-3 (11). PARP-1 has been identified as a nucleolar protein in mammalian cells (17). TcPARP, however, is concentrated in the nucleus upon stimuli such as DNA damage. As it occurs with PARP from other species, DNA breaks induce TcPARP activation, leading to nuclear PAR production (13).

In early divergent unicellular eukaryotes, such as the members of the Trypanosomatidae family, mechanisms involved in the nucleolar formation at the end of closed mitosis are poorly understood. Nucleologenesis occurs as a continuous process that does not require the concentration of nucleolar material within intermediate nuclear bodies such as prenucleolar bodies, suggesting the direct transmission of preassembled nucleolar complexes to daughter nuclei at the end of mitosis. Interestingly, in *T. cruzi* the presence of a thread connecting the two dividing nuclei has been reported (18).

In this work, we identify the N-terminal domain as being sufficient for TcPARP nuclear localization and describe for the first time that TcPARP is enriched in the parasite nucleolus. We also report that both TcPARP and PAR are present in a thread that connects two dividing nuclei and co-localize with nucleolar proteins. These results could extend PAR role well beyond the largely described DNA damage response protein in trypanosomatids.

## Materials and methods

### Parasite cultures

*Trypanosoma cruzi* epimastigotes (CL Brener strain) were cultured at 28 °C in liver infusion tryptose (LIT) medium (5 g.L^-1^ liver infusion, 5 g.L^-1^ bacto-tryptose, 68 mM NaCl, 5.3 mM KCl, 22 mM Na_2_HPO_4_, 0.2% (w/v) glucose and 0.002% (w/v) hemin) supplemented with 10% (v/v) fetal bovine serum, 100 U.mL^-1^ penicillin and 100 mg.L^-1^ streptomycin.

### Plasmid constructions

Several sets of primers were used for the amplification of different TcPARP protein domain combinations by PCR using a plasmid bearing a copy of *T. cruzi* PARP gene (GenBank: DQ061295) as a template (11). PCR products were cloned into a pGEM-T Easy vector and sub-cloned into a pRIBOTEX vector to overexpress the different TcPARP constructs tagged with HA. TcPARP full-length was cloned into pTEX-GFP vector. *T. cruzi* epimastigotes of the CL Brener strain were transfected with the above-mentioned construct as previously described (13). Stable cell lines were achieved after 60 days of treatment with 500 μg mL^-1^ of G418. Full-length TcPARP encompasses from amino acid 1 to 592; ΔN from 127 to 592; ΔNW from 212 to 592; ΔRC from 1 to 211; ΔWRC from 1 to 126 and ΔNRC from 127 to 211. The recombinant proteins were overexpressed in *T. cruzi* epimastigotes and the presence of chimeric proteins was confirmed by Western Blot (S1A Fig). Growth curves of transgenic parasites showed no significant differences compared to wild type epimastigotes (S1B and S1C Figs).

### SDS–PAGE and Western blot analysis

Whole cell lysate of transgenic epimastigotes was quantified using the Bradford method. Proteins (40 µg) were run on 10% SDS–PAGE gel and transferred to an Amersham Hybond-ECL nitrocellulose membrane (GE healthcare), according to the manufacturer’s instructions. Immunodetection of TcPARP constructs was carried out using a 1:1000 dilution of rat polyclonal antibody directed against HA, influenza virus hemagglutinin (Roche), followed by 1:4000 anti-rat horseradish peroxidase (HRP) conjugated antibody. For full-length TcPARP-GFP, anti GFP (B-2): sc-9996 - Santa Cruz Biotechnology antibody was used. The signal was detected with the Western Lightning Plus-ECL kit (PerkinElmer).

### Immunofluorescence

For immunolocalization of TcPARP, transgenic *T. cruzi* epimastigotes (10^7^ parasites ml^-1^) were treated with 300 µM H_2_O_2_ for 10 min, fixed for 20 min with 4% (w/v) formaldehyde in PBS, permeabilized with fresh PBS – 0.1% Triton X-100 and blocked for 1 h at room temperature with 3% (w/v) BSA in PBS. For full-length TcPARP, an anti-GFP antibody 1:500 was used and TcPARP constructs were detected with 1:500 rat polyclonal antibody directed against HA, followed by 1:500 Alexa Fluor 488 goat anti-rat IgG antibody (Sigma-Aldrich) conjugated antibody. The nucleolus was identified using 1:100 L1C6 monoclonal antibody and mouse monoclonal anti α-Tubulin (Sigma-Aldrich) for α-Tubulin. Alexa Fluor 594 goat anti-mouse IgG (Sigma-Aldrich) conjugated antibody was used as a secondary antibody. PAR binding reagent was rabbit polyclonal anti PAR (BD Pharmingen™). Nuclei were stained with 2 µg ml^-1^ DAPI (Sigma) in PBS. Coverslips were mounted with VectaShield® and then visualized using an Olympus BX41/FV300 or Leica SPE confocal microscope. Control images, without primary antibodies, were taken in an analogous condition, during the same microscopy session.

## Results

### TcPARP is enriched in the nucleolus of epimastigotes

As already known from previous research, subcellular localization of TcPARP is altered in response to genotoxic stress, which stimulates TcPARP to re-localize and accumulate in the nucleus in the epimastigote form of the parasite (13). Constructs with different arrangements of TcPARP domains were designed and the recombinant proteins were overexpressed in *T. cruzi* epimastigotes. Actively growing cells were examined for the distribution of the recombinant proteins after treatment with 300 µM H_2_O_2_ for 10 minutes, which has been shown to induce nuclear translocation of endogenous TcPARP (11). Nuclear localization was observed for full length TcPARP and the constructs bearing the N-terminal domain, under control conditions or after H_2_O_2_ treatment. However, TcPARP constructs which lacked the N-terminal domain did not localize to this organelle even after genotoxic stress, showing a scattered pattern throughout the cytoplasm (S2). This result indicates that the N-terminal domain is necessary for the nuclear localization of the protein.

Notoriously, nucleolar enrichment of the recombinant proteins that bear the N-terminal domain (TcPARPΔWRC, TcPARPΔRC and TcPARP-FL) in overexpressing epimastigotes could be observed (Fig 1). A higher intranuclear fluorescence density of the anti-tag signal was detected within the area with reduced DAPI staining. This region corresponds to the nucleolus, which has low DNA content. In order to confirm this localization in transgenic epimastigotes, a co-localization assay was carried out using the nucleolus marker L1C6, which recognizes a nucleolar antigen both in *T. brucei* and *T. cruzi* (19). Figure 1A shows the location of L1C6 in the nucleolar structure of CL Brener wild type epimastigotes. The signal corresponding to TcPARP-FL overlaps with the labeling of L1C6 inside the nucleus of the parasites, which asserts the sub-nuclear localization of this protein. The same phenomenon could be observed for truncated TcPARP constructs, including the N-terminal domain, where the fluorescent signal co-localizes with nucleolus marker L1C6 (Fig 1B). We noticed that in some parasites, both TcPARP (full length or truncated versions) and L1C6 nucleolar marker were also found in the nucleoplasm. In *Drosophila* it has been reported that PARP-1 depletion resulted in a mis-localization of nucleolar specific proteins and loss of nucleolar structure (17). To determine whether TcPARP enzymatic activity and the presence of PAR are essentials for maintaining nucleolar integrity in *T. cruzi*, we inhibited TcPARP activity by adding the NAD^+^ analogue, 3 aminobenzamide (3AB) or the more selective TcPARP inhibitor, Olaparib (20). We did not detect delocalization of the nucleolar marker in the presence PARP inhibitors, ruling out this hypothesis (S3 Fig). Other possibility would be that the high transcriptional activity generates the delocalization of nucleolar markers. To test this hypothesis, we overexpressed a non-nuclear protein such as PI3K TcVps34 in epimastigotes, both in absence or in the presence of cycloheximide. We did not observe differences in the overexpressing parasites in comparison to the control, except for a slight decrease in nucleolar marker in cycloheximide treated parasites. The dispersion of L1C6 could therefore be attributed to epimastigotes in stationary phase as described by Gluenz et al. (21).

**Fig 1.**
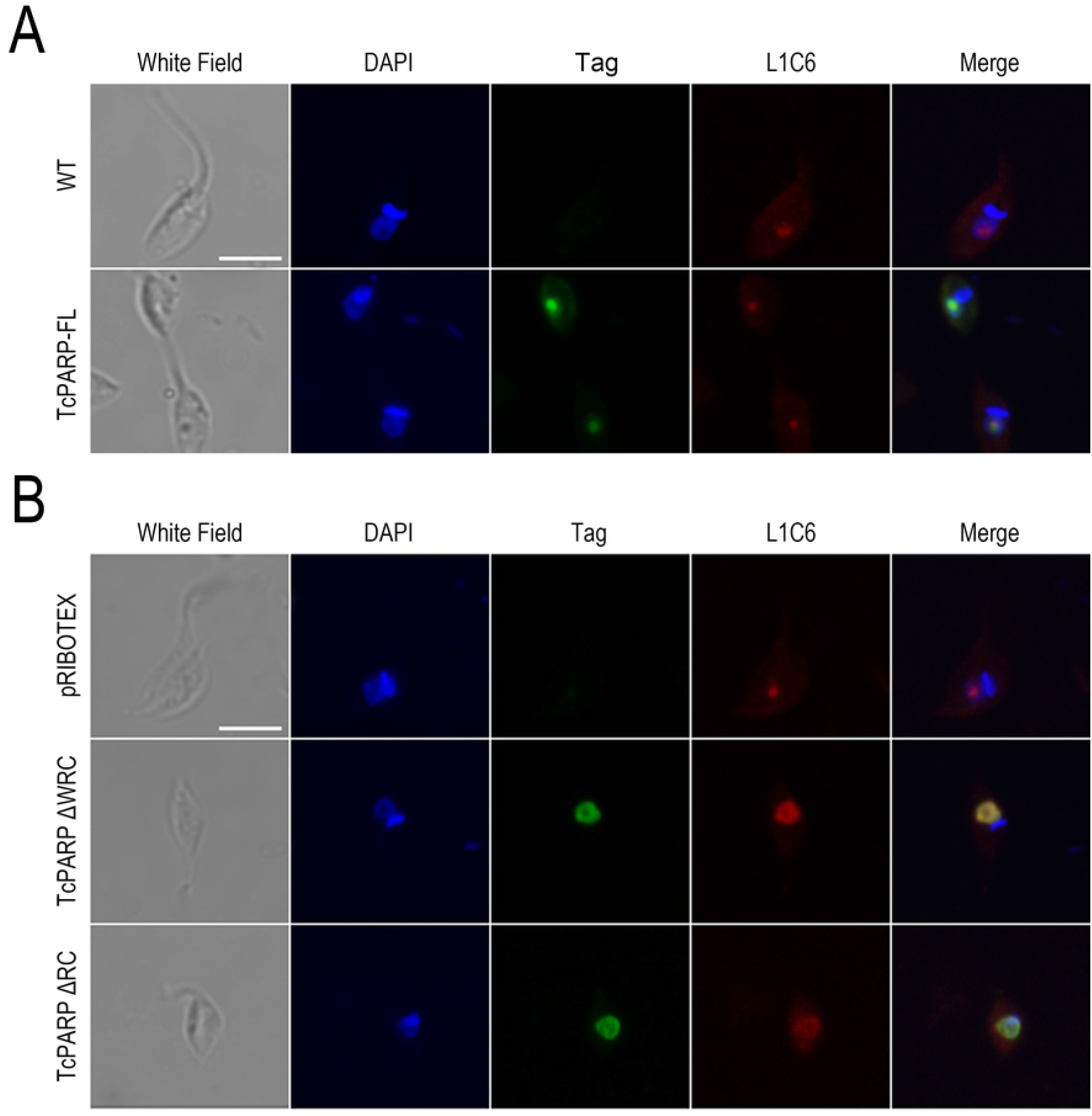
Co-localization of TcPARP constructs with nucleolus marker L1C6. A) Indirect immunofluorescence (IF) of Wild Type or TcPARP-FL overexpressing epimastigotes. B) IF of epimastigotes overexpressing the empty vector pRIBOTEX, TcPARP ΔWRC or TcPARP ΔRC. In both cases, detection of nucleolar region was performed using an antibody directed against L1C6 and the TcPARP construct with the corresponding anti-tag antibody. Bar: 5 μm.

### TcPARP is present in a connecting wire between nuclei of dividing epimastigotes

*T. cruzi* epimastigotes undergoing the cellular division process show the presence of a thread connecting the two dividing nuclei (18), which is associated to nucleolar proteins. We confirmed co-localization of TcPARP with nucleolar proteins in this narrow link between both nuclei by indirect immunofluorescence. The presence of L1C6 could be identified in the entire structure of the wire between cores, co-localizing with the truncated TcPARPΔWRC version of TcPARP or the complete protein (Fig 2). Co-localization of TcPARP with tubulin in the connecting wire is shown in Fig 3, which implies the presence of TcPARP in the mitotic spindle during nuclear segregation and brings forward possible new functions for TcPARP that need further study.

**Fig 2.**
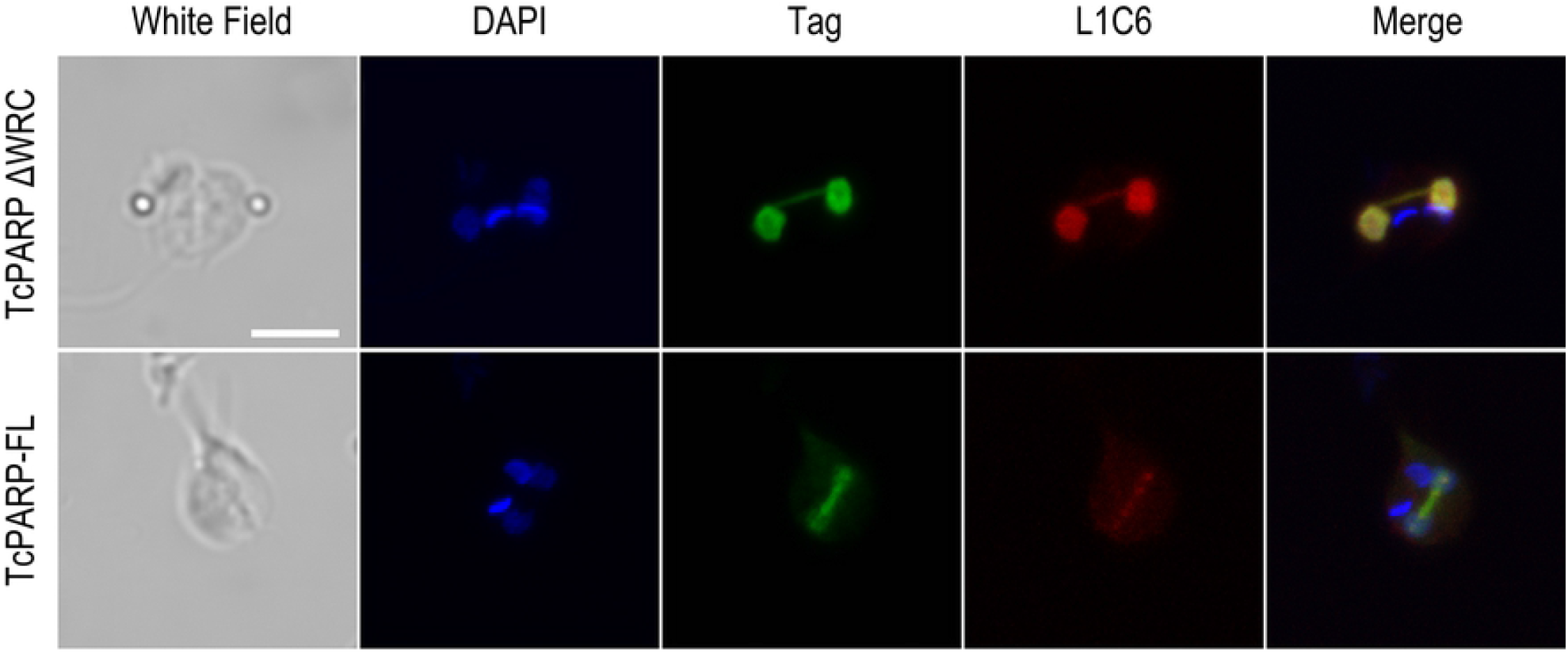
Co-localization of TcPARP with nucleolar material in the connecting wire. Indirect immunofluorescence of epimastigotes overexpressing TcPARP-FL or the N-terminal domain of TcPARP (TcPARP ΔWRC). Nucleolar proteins were evidenced with antibody directed against L1C6. Recombinant TcPARP-FL and chimeric peptides were detected with the correspondent anti-tag antibody. Bar: 5 μm.

**Fig 3.**
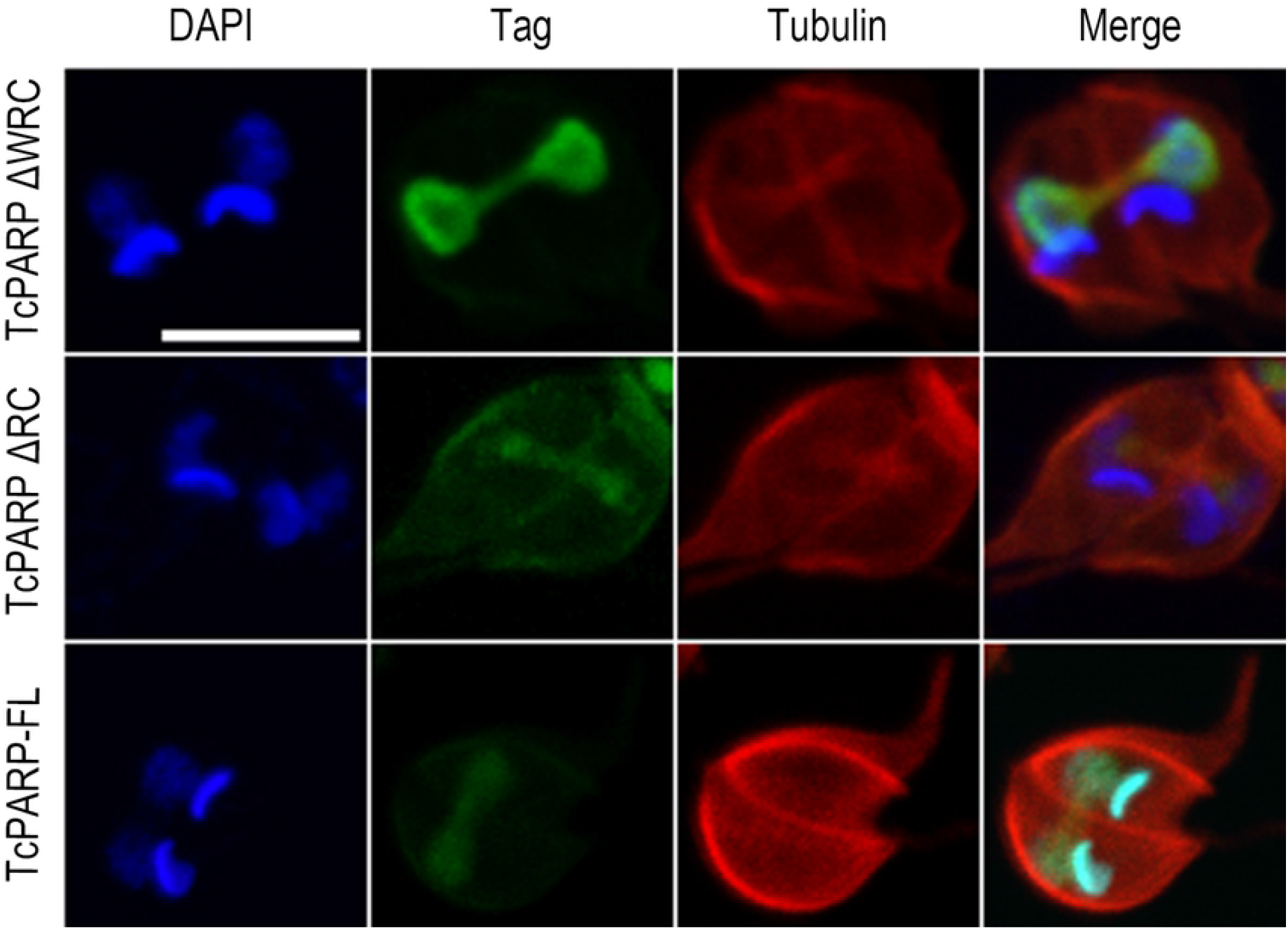
Co-localization of TcPARP in connecting wire with tubulin thread. Indirect immunofluorescence of epimastigotes overexpressing TcPARP-FL or the N-terminal domain of TcPARP (TcPARP ΔWRC and TcPARP ΔRC). Tubulin was detected by an antibody directed against α-Tubulin and recombinant variants of TcPARP with the correspondent anti-tag antibody. Bar: 5 μm.

### PARylated proteins are present in the connecting thread

We investigated if TcPARP was catalytically active in the thread. The presence of PAR was evaluated in epimastigotes that overexpressed the complete TcPARP or the domains responsible for its nuclear localization. Basal levels of PAR are low and therefore hardly detectable by the methods commonly employed (11), therefore, we subjected the parasites to a genotoxic stimulus in order to enhance PAR production. Figure 4 shows the presence of PAR in the nucleus of H_2_O_2_-treated parasites; PAR production co-localizing with the TcPARP constructs is highlighted in the thread that connects the dividing epimastigotes. PAR localization showed the same pattern observed in epimastigotes expressing the empty vector pRIBOTEX, or in wild type parasites (Fig 4). This demonstrates that PARylated proteins are present in this structure as a consequence of endogenous TcPARP activity.

**Fig 4.**
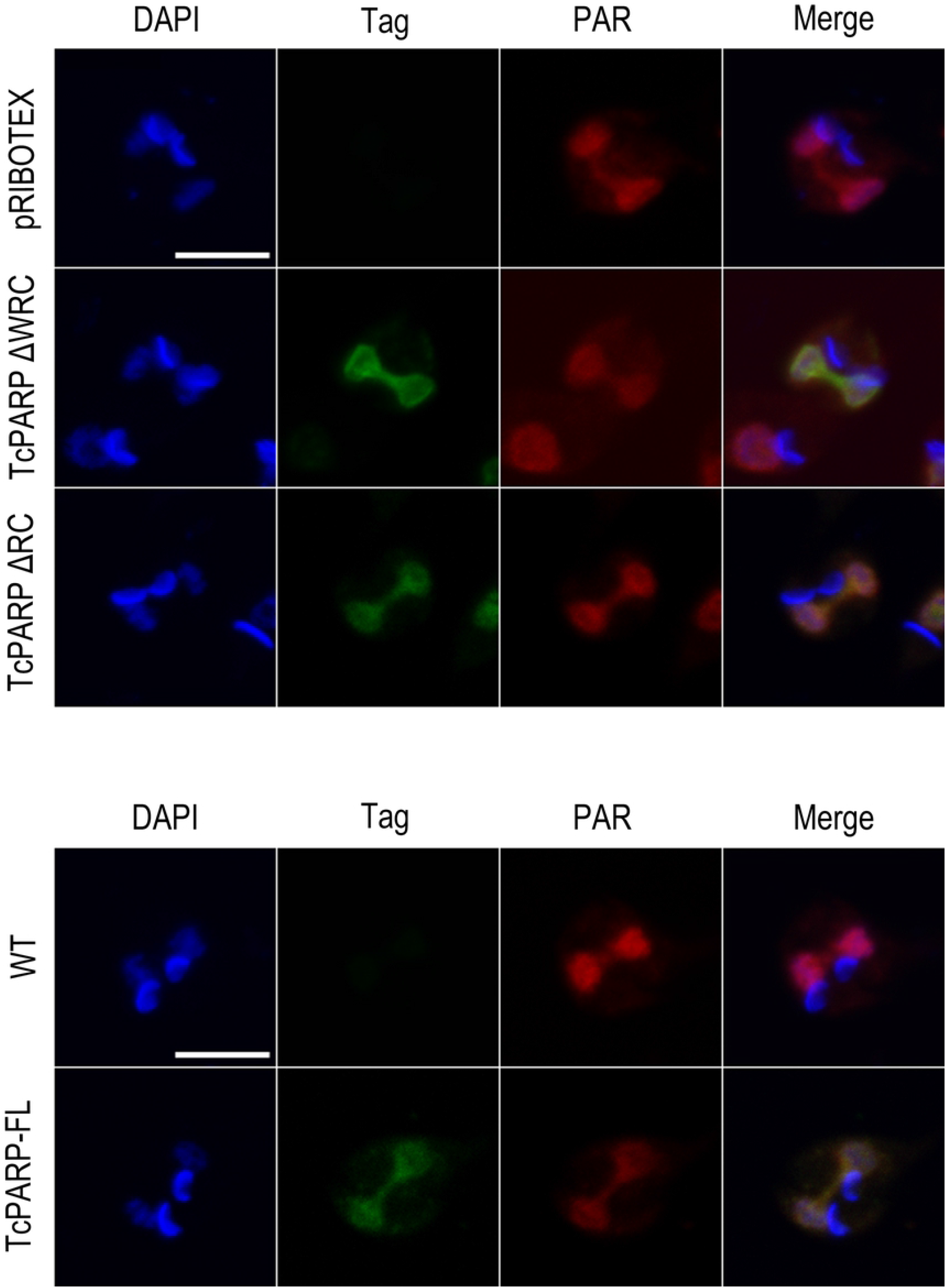
Presence of PAR in the connecting wire between nuclei of dividing epimastigotes. Upper panel: Indirect immunofluorescence (IF) of epimastigotes bearing the empty vector pRIBOTEX or overexpressing the TcPARPΔWRC or TcPARPΔRC constructs, after 300 μM H_2_O_2_ treatment for 10 minutes. Lower panel: IF of wild type epimastigotes or TcPARP-FL overexpressing parasites after 300 μM H_2_O_2_ treatment for 10 minutes. In both cases, PAR was detected using with a polyclonal antibody directed against PAR chains and recombinant variants of TcPARP with the correspondent anti-tag antibody. Bar: 5 μm.

## Discussion

In *Trypanosoma cruzi*, TcPARP translocates to the nucleus under genotoxic stress and in this work we demonstrate that the N-terminal domain is sufficient for this translocation. Despite that no canonical NLS has been found, roughly 30% of N-terminal amino acids are basic (13), which indicates the possible presence of a nuclear localization signal within this region. Interestingly, the TcPARP N-terminal contains six repeats of the amino acid sequence AAKKA, which is present in *T. cruzi* histone H2A and in all H1, which could mediate the DNA binding necessary for TcPARP functions. A *T. cruzi* importin α with the classical structure of ARM repeats, was recently reported (22). This protein was functional as a nuclear transport factor in these parasites, supporting the idea that this import machinery was an early acquisition of the eukaryotic cell. Numerous authors have described the sub-nuclear localization of PARP-1 in *Drosophila* and in mammalian cells (17,23–27), where up to 40% of PARP-1 is enriched in the nucleolus. Our finding that TcPARP localizes in the nucleolus of *T. cruzi* epimastigotes is consistent with that data. Altogether, our results demonstrate that the N-terminal domain of TcPARP is involved in nuclear and nucleolar targeting.

In *Trypanosoma cruzi* closed mitosis occurs, in which the chromatin does not condense and the nucleolus is not clearly identified. The nucleolar structure persists throughout closed mitosis, where the nucleolus is elongated and, finally, divided into two structures (28). Observation of trypanosomes expressing various fusion proteins allowed visualization of mitotic cells connected by a thin thread (29). In dividing cells, we found that TcPARP was located in the connecting wire that separates the daughter nuclei, co-localizing with tubulin and the nucleolar marker. As determined by DAPI staining, nuclear DNA seems to be excluded from the thread indicating a late stage in mitosis. This observation implicates that TcPARP, like other nucleolar components, forms part of the connecting thread after chromosomal segregation. Among the PARP functions in mammalian nucleolus, Raemaekers (2003) identified the modification of NuSAP1 (nucleolar protein associated with spindle 1), a protein involved in the mitotic spindle formation, which concentrates in the nucleolus. In metaphase and early anaphase, NuSAP is redistributed, co-localizing with and stabilizing the spindle microtubules (30). Both NuSAP and tubulin are targets of PARylation (31). Similarly, NUMA1 (nuclear mitotic apparatus protein 1), which participates in the clustering of microtubules into poles as a prerequisite for bipolar spindle organization, is associated with PAR. Its PARylation and PAR-binding property may promote correct assembly of bipolar spindles by crosslinking NUMA1 molecules between spindle poles (32). Thus, the PARylation of spindle proteins associated with microtubules plays an important role in the assembly and function of the mitotic spindle, where PARP could work as a Microtubules Associated Protein. In addition to this function, PAR has been postulated as a structural component of the spindle that contributes to creating the force exerted by microtubules to elongate the spindle and separate chromosomes (33). Furthermore, ECT2 (epithelial cell trans-forming sequence 2 oncogene), necessary for the control of cytokinesis, is recruited by PARylated α-tubulin to the spindle during metaphase as a prerequisite for functional cytokinesis and completion of mitosis (34). Then, the nucleolar location of TcPARP and associated PARylated proteins during mitosis could be involved in spindle formation, not only through the PARylation of nucleolar proteins but also as a result of their structural functions. The possible participation of TcPARP in the nucleolus as an enzyme related to the formation of the mitotic spindle requires deeper investigation. Thus, TcPARP could be performing a wide array of new roles, leaving aside the notion that this enzyme is functional only in the nucleus.

## Acknowledgments

The authors are grateful to Dr. Daniel Sanchez, Instituto de Investigaciones Biotecnológicas (IIB), Universidad Nacional de San Martín – CONICET, for providing L1C6 antibody. We are also grateful to Dr. Alejandra Schoijet, Instituto de Investigaciones en Ingeniería Genética y Biología Molecular “Dr. Héctor N. Torres”, for the PI3K TcVps34 epimastigotes.

## Supporting information

**S1A Fig. Constructs bearing different combinations of TcPARP domains expressed in transgenic epimastigotes**. Left Panel: Schematic Diagram of truncated TcPARP recombinant proteins made through the combination of different domains and fused to HA tag. TcPARP full length was tagged with GFP. Numbers under each diagram indicate amino acid positions. Theoretical molecular weight was calculated considering the molecular weight of HA (1 kDa) or GFP (27 kDa). Right panel: Western Blot on total extract of *T. cruzi* epimastigotes transfected with plasmids bearing the indicated constructs, using the appropriate antibody against the tag. The arrows indicate the expected molecular weigth corresponding to the expression of the different constructs.

**S1B Fig. Parasite growth curves**. Cultures of CL Brener strain-epimastigotes in the exponential growth phase were collected by centrifugation at 3000g/5min at room temperature, washed with PBS and suspended in LIT (6.10^6^ parasites ml^-1^). Parasite growth was measured daily by 600 nm absorbance readings in 96-wellmicroplates.

**S1B Fig. Relative growth**. Abs_600nm_ at day 4 was normalized to the initial value (Abs t_0)_.

**S2 Fig. Subcellular location of TcPARP constructs**. Indirect immunofluorescence of *T. cruzi* epimastigotes (CL Brener strain) that overexpress TcPARP-FL or different combinations of protein domains, under basal conditions (Control) or treated with 300 μM hydrogen peroxide (H_2_O_2_) for 10 minutes. (A) TcPARP-FL. (B) TcPARPΔN. (C) TcPARPΔNW. (D) TcPARPΔRC. (E) TcPARPΔWRC. (F) TcPARPΔNRC. Bar: 10 μm

**S3 Fig. Dispersion of the nucleolus**. Actively growing wild type epimastigotes preincubated for 1 h in the presence of the NAD^+^ analogue, 3 aminobenzamide (3AB), or the TcPARP inhibitor, Olaparib; and PI3K TcVps34 overexpressing parasites (Overexpressants) in the presence or absence of cycloheximide, were fixed and labeled with L1C6 antibody. Bar: 5 μm

